# Feedback Control of Neuronal Excitability and Epileptiform Bursting using a Photocaged Adenosine A_1_ Agonist

**DOI:** 10.1101/2024.06.17.597004

**Authors:** Erine Craey, Serge Van Calenbergh, Jeroen Spanoghe, Marijke Vergaelen, Lars E. Larsen, Evelien Carrette, Jean Delbeke, Kristl Vonck, Paul Boon, Wytse J. Wadman, Robrecht Raedt

## Abstract

Adenosine is a potent regulator of neurotransmission and neuronal excitability through activation of G_i_ protein-coupled adenosine A_1_ receptors (A_1_Rs). It has gained interest as a potential anticonvulsant due to its endogenous involvement in ending ongoing seizure activity.

A recently developed coumarin-caged derivative of the A_1_R agonist N^6^-cyclopentyl-adenosine (CPA), cCPA, was used for photo-uncaging of CPA with millisecond flashes of 405 nm light. At population level, CPA reduces Schaffer Collateral stimulated extracellular dendritic field potentials (FPs) in the CA1 region of the hippocampus with an ED_50_ of 44.1±2.8 nM and a Hill coefficient of 3. Response onset is CPA dependent and takes less than seconds, while recovery is CPA independent with a time constant of around 20 minutes.

A closed-loop feedback system used the amplitude of evoked dendritic FPs to photorelease CPA and was able to control FP amplitude to user defined levels between 10% and 90% of baseline level. In the acute elevated potassium model of epilepsy raising extracellular K^+^ to 8.5 mM enhances neuronal excitability and induces regularly occurring epileptiform bursts, but FPs evoked with low intensity could still continuously monitor excitability without interfering with bursting.

In this model the closed-loop system that controlled CPA release, was able to suppress epileptiform bursting, while maintaining an acceptable level of functional neurotransmission. Including in the control algorithm a second parameter that combined population spike amplitude and number of population spikes, enabled the system to automatically find a level of functional neurotransmission that was just below the threshold for multiple spiking and epileptiform bursting.

The combination of photopharmacological adenosinergic modulation with real-time FP monitoring provides a first step towards closed-loop precision treatment for diseases related to neuronal hyperexcitability such as epilepsy.

## 1. Introduction

Adenosine is a potent modulator of neuronal excitability and it might even act as an endogenous anticonvulsant because its increase during seizures counteracts ongoing seizure activity [1]. These effects are mainly mediated by adenosine A_1_ receptors (A_1_Rs), which are G_i_ protein-coupled receptors. Activation of the A_1_R initiates nucleotide exchange on the heterotrimeric G_i_ protein (GDP to GTP) leading to dissociation into G_iα_ and G_βγ_ subunits, which in turn interact with their downstream effectors. The G_βγ_ subunits interact with voltage-gated calcium channels (VGCC), inhibits their opening in response to depolarization, so reducing calcium influx and neurotransmitter release. The G_βγ_ subunits also bind to G protein-mediated inwardly rectifying potassium (GIRK) channels; the resulting increase in K^+^-conductance leads to hyperpolarization and reduces neuronal firing. The intrinsic GTPase activity of G_iα_ will hydrolyse its bound GTP to GDP, whereafter G_iα_ will reassociate with the G_βγ_ subunit. The rate of GTP hydrolysis is slow and independent of receptor occupancy [2]–[4].

The potent and long-lasting effect of A_1_R activation in neurons has gained attention as a potential target for epilepsy treatment [5]–[8]. However, the development of adenosinergic therapeutics for epilepsy has been hindered by side effects associated with activation of these receptors ubiquitously present throughout the body [9]–[11]. Most pronounced are cardiovascular adverse effects such as decreased arterial blood pressure and heart rate [12], [13]. To minimize unwanted systemic side effects, focal adenosinergic treatment is preferable which selectively suppresses neuronal excitability in brain regions associated with seizure generation.

Recently a photocaged derivative of the A_1_R agonist N^6^-cyclopentyladenosine (CPA) was synthetized by conjugating its 6-amino group to 7-(diethylamino)-4-(hydroxymethyl)coumarin (DEACM) [14]. The resulting caged CPA (cCPA) had a more than 1000-fold decreased affinity for the A_1_R and was irreversibly released by 405 nm light. The inactivity of cCPA at a concentration of 3 µM was confirmed in acute hippocampal brain slices where neurotransmission was assessed by electrical stimulation of the Schaffer Collaterals which evoked extracellular field potentials (FPs) in the CA1 region [14]. Micromolar concentrations of cCPA did not affect the FPs while flashes of 405 nm light resulted in fast decrease and slow recovery of FP amplitude and thus induced a transient reduction of neurotransmission. Based on these observations an on-off closed-loop control paradigm was implemented, where repeatedly evoked FPs were monitored and a hardware system delivered light flashes that released CPA on pre-set level crossing [14]. The system was able to stabilize the FP at an amplitude that was 50% of its baseline value. This led to the hypothesis that it might be possible to use adenosine-based photopharmacology for controlling the level of excitatory neurotransmission within limits that in pathological conditions like epilepsy prevent seizure activity while ensuring a healthy level of functional neurotransmission.

In the current study a multi-electrode array (MEA) was chosen because it allowed recording of dendritic as well as somatic FPs in acute rat hippocampal slices in response to Schaffer Collateral stimulation in region CA1. In this preparation the dynamics of A_1_R-mediated inhibition of neurotransmission and neuronal excitability with CPA were quantified at the population level. Pulsed photorelease of CPA demonstrated the reduction in excitability that is induced by A_1_R activation, while the use of the highly specific competitive antagonist DPCPX proved that these effects are indeed mediated by the A_1_Rs.

An extended version of the previously described closed-loop feedback system analysed FPs on two MEA electrodes in real time and fed the resulted parameters into an algorithm that defined the moment and duration of the CPA releasing light flashes. As the ultimate goal of the system was suppression of epileptiform bursting while still permitting neurotransmission at an acceptable level, an acute (slice) model with epileptiform bursts was implemented by elevating extracellular potassium concentration from 3.25 to 8.5 mM. This model is characterized by enhanced excitability and regular occurring epileptiform bursts. It was demonstrated that despite the bursting, FPs can still be used to monitor excitability and as input to the feedback system. The latter was able to reduce excitability and suppress bursting while still permitting neurotransmission at a pre-set level. The advanced feedback system allows fine tuning of the control loop in much greater detail than was investigated here, but the experiments provide a clear proof of concept of this approach.

## 2. Materials and Methods

### Hippocampal slice preparation

Thirty-seven male Wistar rats of 4-6 weeks old (Janvier Labs, France) were housed with a 12/12-hour light dark cycle and received food and water *ad libitum*. Animals were treated according to the European guidelines (directive 2010/63/EU) and experimental procedures were approved by the Animal Experimental Ethical Committee of Ghent University (ECD 19/29). Rats were killed by decapitation under isoflurane anaesthesia. The brain was quickly removed and placed in ice-cold carbogen-saturated (95% O_2_/5% CO_2_) artificial cerebrospinal fluid (aCSF) of the following composition (mM): NaCl 124, NaHCO_3_ 26, KCl 2, KH_2_PO_4_ 1.25, CaCl_2_ 2, MgSO_4_ 2 and glucose 10 (pH=7.4). Transverse slices of 350 *µ*m thickness were prepared from the ventral hippocampus using a VT1200S microtome (Leica Microsystems, Wetzlar, Germany) and stored in carbogen-saturated aCSF at room temperature for at least one hour until use. In the elevated potassium experiments, extracellular potassium concentration ([K^+^]_o_) was raised to 6.5 or 8.5 mM by the addition of the required amount of KCl from a 1 M stock.

### Electrophysiological recordings

Slices were transferred to the MEA-chamber (MultiChannel Systems (MCS), Reutlingen, Germany) and continuously superfused with carbogen-saturated aCSF (flow rate: 2 mL/min) that was preheated to 34^◦^C by a cannula (PH01, MCS) positioned at the inlet. This maintained bath temperature at 31^◦^C. A custom-made nylon grid ensured slice position and contact with the electrodes. Extracellular recordings were performed with a planar MEA (MCS) containing a square grid of 60 titanium nitride coated electrodes (30 *µ*m diameter; 200 *µ*m spacing), one of which was used as the common reference. Data were acquired with a high-bandwidth MEA1060-BC preamplifier and digitized at either 10 or 25 kHz with 16-bit resolution, using MC_Rack software (Version 4.6.2, MCS).

After an equilibration period of at least 15 minutes, slice quality was evaluated by applying a monophasic negative voltage stimulus (duration 100 *µ*s; amplitude 2000 mV) from an external stimulus generator (STG4002, MCS) to the user-selected MEA electrode located closest to the Schaffer Collaterals. Stimulation parameters and experimental timing were controlled using MC_Stimulus software (MCS). Only slices where at least three channels within the CA1 region displayed an extracellular postsynaptic potential (EPSP) with an amplitude larger than 150 *µ*V were included for further analysis. In all experiments, FPs were repeatedly evoked with an interval of 10 seconds. Paired pulse stimuli with identical amplitude and an interstimulus interval of 20 ms were often used. In certain experiments single and paired pulse stimulations were alternatingly given. A stimulus intensity (I_xx_) is defined as the intensity that evoked an EPSP amplitude that is xx% of the largest response. Flash photolysis of cCPA was induced with computer-controlled light flashes (25, 50 or, 500 ms) delivered by a 405 nm light-emitting diode (LED, M405FP1, Thorlabs, Bergkirchen, Germany), positioned above the MEA chamber. Light energy output was measured with a photodiode power sensor (S120VC, Thorlabs, Bergkirchen, Germany), placed at the MEA chamber position, and set at 4.0 mW. Experiments with cCPA were always performed in the dark.

In the simple closed-loop experiments light flashes were triggered by MCS built-in hardware that detected level crossing within a time window of 2 to 19 ms after the stimulus in a user-selected MEA channel. This generated a digital pulse of fixed duration that drove the LED. In the more complex closed-loop experiments during elevated potassium conditions, an external ADC (NI-USB-6259, National Instruments, Austin, Tx, USA) sampled two user-selected MEA channels, routed via the audio channels, that were analysed in real-time with the same MATLAB software as in the offline analysis. A user-defined algorithm, that used the parameters EPSP amplitude and coastline index (CLI), decided if, when, and for how long the LED light flash had to be given via the DAC output of the same multipurpose DAQ.

### Data Analysis

Offline analysis of the FPs was performed with custom-made software written in MATLAB (MathWorks, Natick, MA, USA). FPs were offset corrected using 5 ms of the signal before the first stimulus, the stimulus artefact was suppressed, and the signals were low-pass filtered at 1200 Hz. The amplitude of the dendritic EPSP was calculated as the difference between the minimal amplitude and the value determined 2 ms before its stimulus (**Figure 1C**). Two other forms of EPSP amplitude estimation were also employed: 1) The minimal/maximal slope of the falling/rising phase of the FP gives a parameter closer to the underlying synaptic current, but it is also much more sensitive to noise and filtering. 2) Assuming that a set of EPSPs have the same shape and only differ by a gain factor, those gain factors can be estimated with the least square scaling to the mean (LSSM) method (described below). All three methods were used to calculate EPSP amplitude in the FP analysis; the slope is only preferred when a large population spike (PS) obscures the peak of the synaptic current while LSSM can only be used in the absence of a PS. For consistency and comparability maximum amplitude is used throughout this paper (unless indicated otherwise) and it was always checked that conclusions were not dependent on the used quantification method. The amplitude of the PS, superimposed on the somatic EPSP, was calculated as the difference between the negative peak of the PS and the tangent connecting the two adjacent positive peaks (**Figure 1C**). The CLI or line length of a trace was calculated as the summed line length of the band-pass filtered (300 and 1200 Hz) FP between 2 and 150 ms after the stimulus.

**Figure 1.**
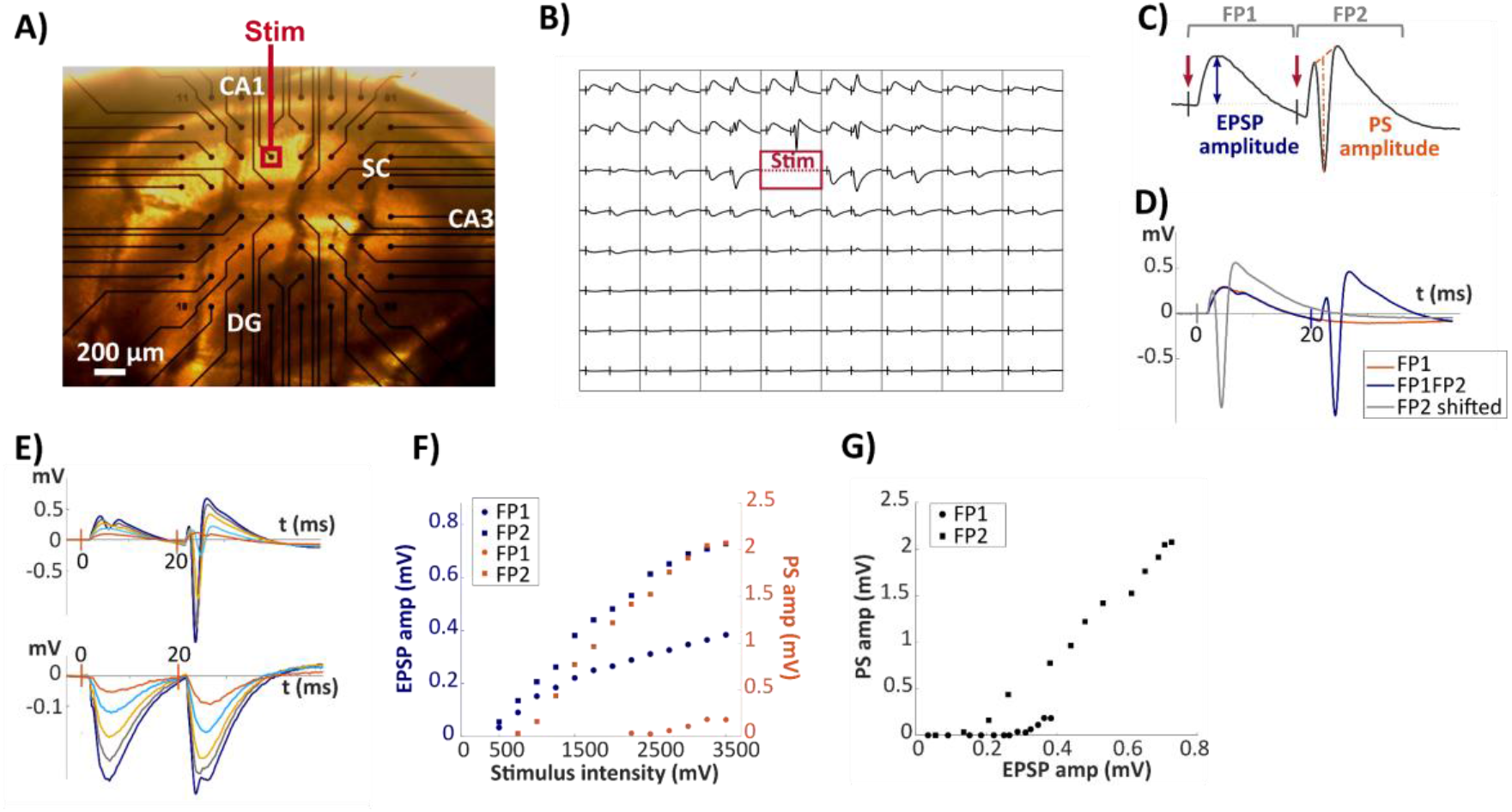
FPs in the CA1 hippocampal region recorded with a MEA. **A)** Transverse hippocampal slice positioned on a 60 channel MEA in such a way that the 8×8 matrix of recording electrodes covered the CA1 region. A user-selected electrode (Stim) delivered electrical stimuli to the Schaffer collaterals (SC). **B)** FPs recorded by the surrounding electrodes demonstrated a negative dendritic EPSP in the Stratum Radiatum and a corresponding positive somatic EPSP in the Stratum Pyramidale, often accompanied by a PS. **C)** Quantification of the two components of the FP: EPSP amplitude (vertical blue double arrow in the response during FP1) and PS amplitude (vertical dotted orange line in the response during FP2). **D)** Overlay of the single pulse response (orange), the paired pulse response (blue) and the subtracted response (grey), shifted leftwards by 20 ms to allow direct comparison of the responses in FP1 and FP2. **E)** Superimposed traces recorded from an electrode in the Stratum Pyramidale (top) and one in Stratum Radiatum (bottom) in an experiment where stimulus intensity was stepwise increased from threshold to saturation to construct a SR-curve. **F)** Peak amplitude of the EPSP (blue) and PS amplitude (orange) from the responses in FP1 and FP2 (**E**) as a function of stimulus intensity. **G)** The responses in (**E**) also allowed to determine the E-curves: PS as a function of EPSP for FP1 (blue) and for FP2 (orange). As the same EPSP gave a substantial larger PS in FP2 compared to FP1, FP2 is considered as the period with higher excitability.

When n experimentally obtained curves only differed by a gain factor g_i_: f_i_(x) = g_i_ * f_mean_(x), i=1:n, the gain factors were estimated with a linear least square fit of the f_i_’s on their mean f_mean_(x). This method is called LSSM and used to estimate the reduction in EPSPs after application of CPA (**Figure 2**). Moreover, it is also used to estimate increase in SR-curves (**Figure 6B**) and to compare the curves that show EPSP amplitude as a function of time (**Figure 6A**). Fitting a specified function, such as the binding function (**eq 1.4**), to a set of data points was done with a nonlinear least square fitting routine provided by MATLAB.

**Figure 2.**
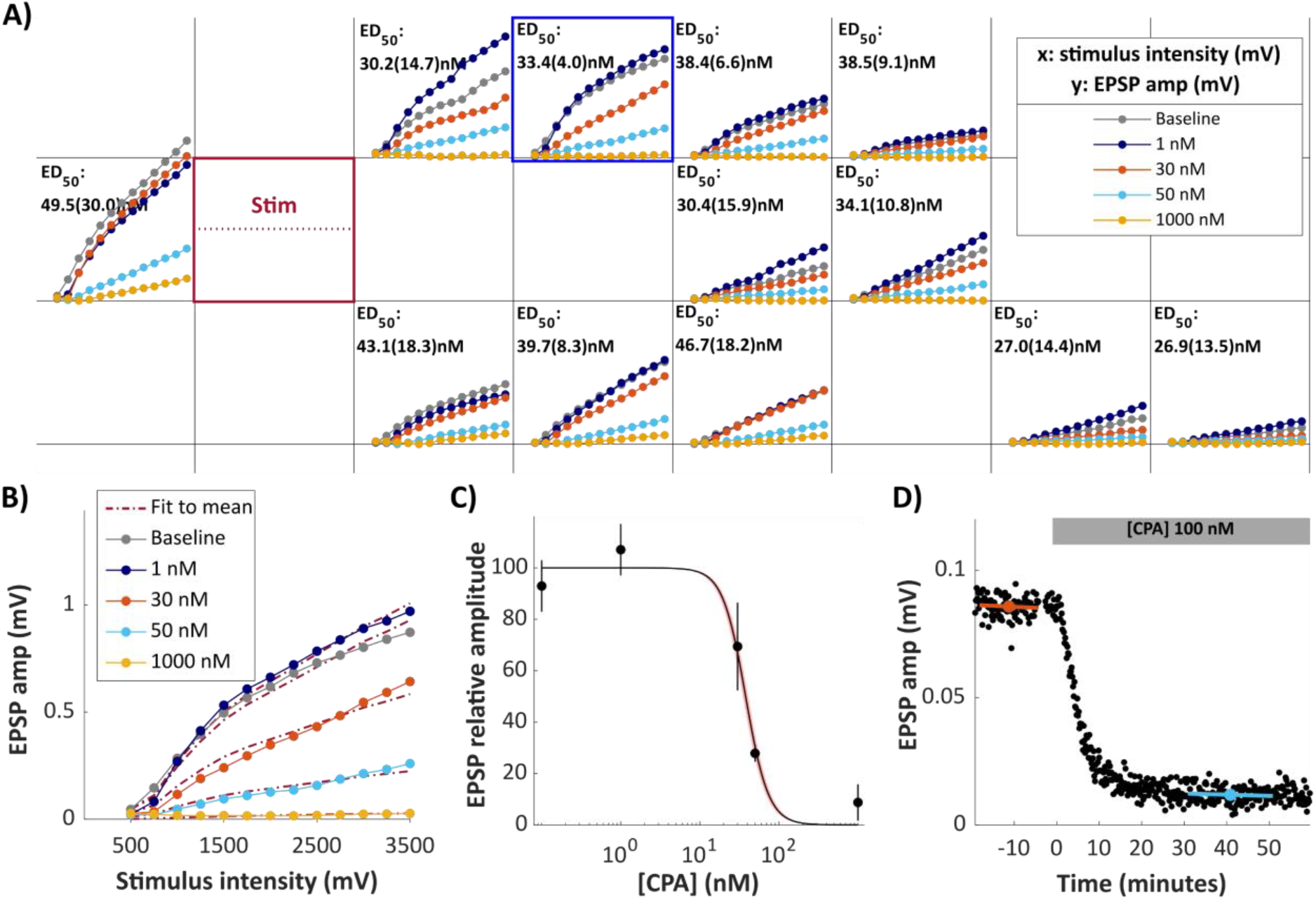
Concentration-dependent reduction of EPSPs by CPA activation of A_1_Rs. **A)** Representative example of the SR relationship (EPSP amplitude as a function of stimulus intensity) measured in the same slice during baseline and after perfusion with four CPA concentrations (1, 30, 50 and, 1000 nM) for electrodes that displayed a relevant EPSP. Stimulus electrode is marked Stim. **B)** SR-curves from the electrode marked with the blue square in (**A**). The red dotted lines indicate the LSSM fits for each curve. **C)** The EPSP amplitude reduction obtained from **B)** as a function of [CPA], normalised to baseline and fitted with a binding equation (**eq 1.4**). In this experiment, a Hill coefficient of 3 and an ED_50_ value of 37.8±1.6 nM were obtained. Data points are weighted averages from the data obtained from 12 MEA channels dispayed in **A)**. Vertical bars indicate the standard deviation. The red shaded area around the curve indicates the region of one standard error of the mean. **D)** Time course of the EPSP amplitude in response to a single CPA concentration change (100 nM), which showed stable suppression in about 10 mins. Assuming a Hill coefficient of 3 and full suppression for high CPA concentrations, the estimated ED_50_ was 54 nM.

To calculate the amount of CPA released per flash two assumptions were made: 1) the flashes are the only source of CPA, and each flash always releases the same amount of CPA which raises [CPA] by Y nM and 2) CPA is removed from the extracellular space by a first order process with a washout time constant of *τ* minutes. During the initiation phase pulses of CPA were given at times t_i_ until the equilibrium was reached at the first pulse after the undershoot: t_n_. At t_n_ the equilibrium concentration CPA_eq_ is described by the following equation:

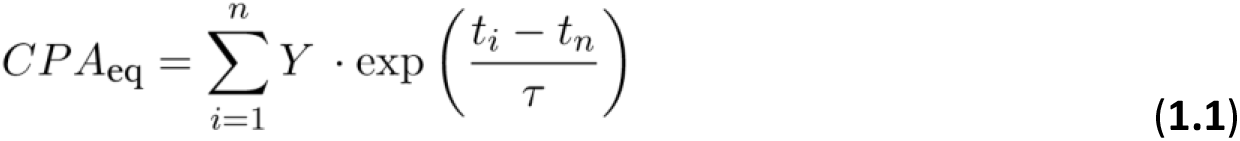

In steady state pulses occurred at times t_s_ between t_n_ and t_end_ keeping CPA_eq_ at the same stable value:

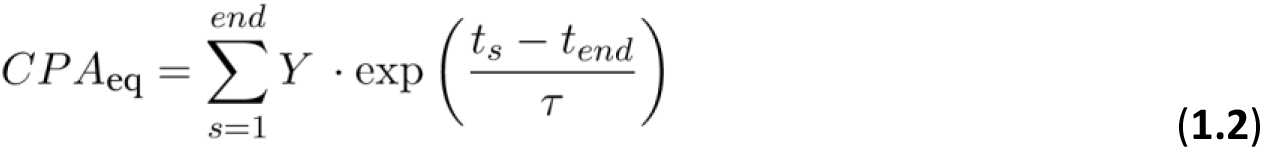

**Eq 1.1** and **1.2** combine into an implicit second order equation that can be numerically solved and leads to two solutions. The one with a realistic washout time constant is used.

### Statistical Analysis

Statistical analysis was performed with Prism (v8.0.2, GraphPad, San Diego, USA) and MATLAB (MathWorks, Natick, USA) software. Data are reported as mean ± standard error of the mean (SEM) with n representing the number of slices, unless indicated otherwise. Means were compared with a Students t-test after normality was validated with the Shapiro-Wilk test. P<0.05 was assumed to reject the null hypothesis.

### Chemicals

CPA was purchased from Tocris Bioscience (Bristol, United Kingdom). DPCPX and DMSO were purchased from Sigma-Aldrich (Missouri, USA). cCPA was synthesized and provided by the Laboratory of Medicinal Chemistry (Ghent University, Belgium)[14]. Stock solutions of CPA (1, 30, 40, 50, 100 and 1000 *µ*M) and of cCPA (3 mM) were made in DMSO and stored at -20^◦^C. The final perfusion solution always contained 0.1% DMSO.

## 3. Results

### 3.1 Evoked field potentials (FPs) monitor neurotransmission and neuronal excitability

A 60-channel MEA was used to record CA1 FPs evoked by electrical stimulation of the Schaffer collaterals in the Stratum Radiatum of rat hippocampal slices (**Figure 1A**). In response to paired pulse stimulation (20 ms interval), on average 18 electrodes in each slice displayed typical CA1 extracellular postsynaptic potentials (EPSPs) (**Figure 1B**). Half of these located in the Stratum Radiatum showed the typical dendritic negative voltage deflection, while the other half located in the Stratum Pyramidale showed the corresponding somatic positive deflection. EPSPs that predominantly reflect (excitatory) glutamatergic synaptic currents were quantified by their amplitude (**Figure 1C**; and Materials and Methods). When stimulus intensity was strong enough, a PS that represents the summed extracellular fields generated by a large number of synchronous action potentials in the pyramidal cell layer was superimposed on the somatic EPSP. The response to the first stimulus of the pair will be denoted as FP1; the one to the second as FP2. In many protocols stimulation was alternated between single and paired pulse, which allowed subtraction of the former from the latter, thus isolating the component induced by the second stimulus (**Figure 1D**).

Each slice experiment started with the determination of the stimulus-response curve (SR-curve) that relates EPSP and PS amplitude to stimulus intensity. To this aim stimuli were applied with stepwise increasing intensities between threshold and saturation (**Figure 1E**) whereafter EPSP and PS amplitude in FP1 and FP2 were plotted as a function of stimulus intensity (**Figure 1F**). The SR-curve for the EPSP quantifies the efficacy of neurotransmission: how much EPSP is obtained for the applied stimulation. The excitability curve (E-curve) is the relation that describes PS as a function of EPSP and is constructed from the SR-curves: how much PS is obtained for a certain amount of EPSP (**Figure 1G**).

Paired pulse facilitation (PPF) was consistently seen in all FPs. PPF implies that the same stimulus intensity leads to a larger EPSP in FP2 than in FP1 (EPSP2/EPSP1>1) and indicates enhanced efficacy of neurotransmission (**Figure 1F**). Despite differences in absolute amplitude, all channels that recorded acceptable signals from the CA1 region in a slice showed a similar value for the PPF. This indicated a considerable redundancy in the information obtained from the various MEA channels. Using the weighted average from all relevant channels avoided a subjective search for the “optimal” channel. LSSM of the SR-curve for EPSP in FP2 to the one in FP1 gave a factor of 1.41±0.03 (*n*=13), remarkably stable over channels and slice experiments. The same EPSP amplitude was associated with a larger PS in FP2 than in FP1, implying that not only neurotransmission was enhanced in FP2 but also that during FP2 neuronal excitability was considerably higher than during FP1 (**Figure 1G**).

### 3.2 The A_1_R agonist CPA modulates synaptic transmission

CPA binding to A_1_R leads to dissociation of G_i_ protein into G_αi_ and G_βγ_. These subunits bind to ion channels where they modulate their biological function: inhibition of presynaptic VGCCs lead to reduction of neurotransmitter release, while coupling to GIRK channels results in membrane hyperpolarization and reduced firing [3]. Essential in this complex biological system is the level of functional relevant G_βγ_ subunit [G], that is increased with a rate (*P_on_*), determined by the occupancy of the A_1_R and it is decreased with rate (*P_off_*), determined by the intrinsic hydrolysis of the subunit and independent of receptor occupancy. The dynamics of [G] in its most simplified form can be described as a first order process:

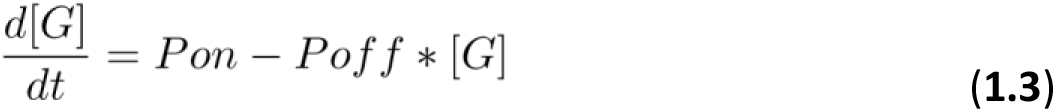

In a previous study the recovery time constant *(1/P_off_)* was determined to be 22±1 mins [14]. Once in steady-state (d[G]/dt = 0), the level [G] is equal to *P_on_/P_off_* and under the assumption of a proportional relation between [G] and the EPSP suppression, the EPSP amplitude in steady state is determined by the binding of CPA to the A_1_R:

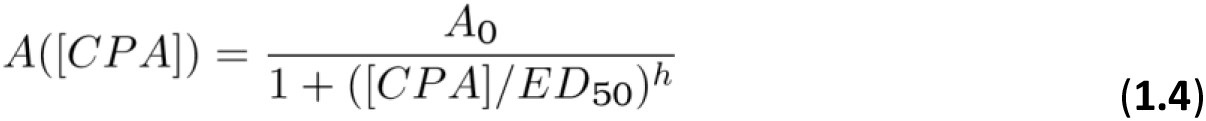

where [CPA] represents the CPA concentration, A([CPA]) is the amplitude of the EPSP in the presence of CPA, *A_0_* the amplitude in the absence of CPA, *ED_50_* the [CPA] value for 50% reduction and *h* the Hill coefficient.

The CPA-concentration dependent EPSP suppression was determined by perfusing the slice with increasing CPA concentrations (1, 30, 50 and 1000nM) in a cumulative sequence starting from baseline. At least 10 minutes after switching the perfusion solution to a higher CPA concentration, three SR-curves were recorded that related EPSP amplitudes to thirteen stimulus intensities (500 to 3500 mV in steps of 250 mV) between threshold and saturation (**Figure 2A**). The SR-curves have a characteristic shape, and LSSM was used to calculate the amplitude reduction (details in Materials and Methods) of all concentration curves (**Figure 2B**; red dotted lines); finally, they were normalized to baseline. EPSP reduction as a function of [CPA] was then fit with the binding equation (**eq 1.4**). Although quite different in amplitude, most electrodes provided similar binding fits. In **Figure 2C** a weighting average of the reduction factors from all relevant electrodes was used to fit the binding curve for this slice. The mean result over all slice experiments gave a Hill coefficient (*h*) of 3 and an ED_50_ of 44.1±2.8 nM (*n=4*). In an additional set of slices, the response to a single concentration of CPA (40 nM (*n=1*), 50nM (*n=2*) or 100 nM (*n*=1)) was tested. Assuming a Hill coefficient of 3 and complete suppression at full receptor occupation, these experiments yielded a comparable ED_50_ of 41.9±6.6 nM. More importantly, they demonstrate that stable levels of suppression are reached (**Figure 2D**).

### 3.3 Changes in hippocampal neurotransmission and excitability after a brief pulse of photoreleased CPA

A previous paper demonstrated the possibility to release CPA from cCPA with short 405nm light flashes resulting in transient suppression of EPSP amplitude [14]. Here we follow up on these experiments to confirm that this transient reduction in EPSP amplitude is indeed mediated by A_1_R activation. Hippocampal slices (*n*=10) were incubated with 3 µM cCPA and exposed to a 500 ms light flash that released CPA (**Figure 3A**). The flash transiently reduced EPSP amplitude to 61.6±4.5% of baseline level. Next, the selective A_1_R antagonist 1,3-dipropyl-8-cyclopentylxanthine (DPCPX) was added to the aCSF in a concentration of 100 nM (*n*=2) or 250 nM (*n*=2). Perfusion with DPCPX increased the EPSP amplitude to 221±60% of baseline for 100 nM and to 225±27% for 250 nM (**Figure 3A**). This increase in EPSP amplitude was, however, not associated with a change in neuronal excitability (**Figure 3B**). Following another 500 ms light flash in the presence of 100 nM or 250 nM DPCPX, the EPSP amplitude was only reduced to 74.4±3% or to 96.5±0.5% respectively. This implied that DPCPX reduced the CPA response by 35% respectively 91% for the two DPCPX concentrations and confirmed that they were A_1_R-mediated.

**Figure 3.**
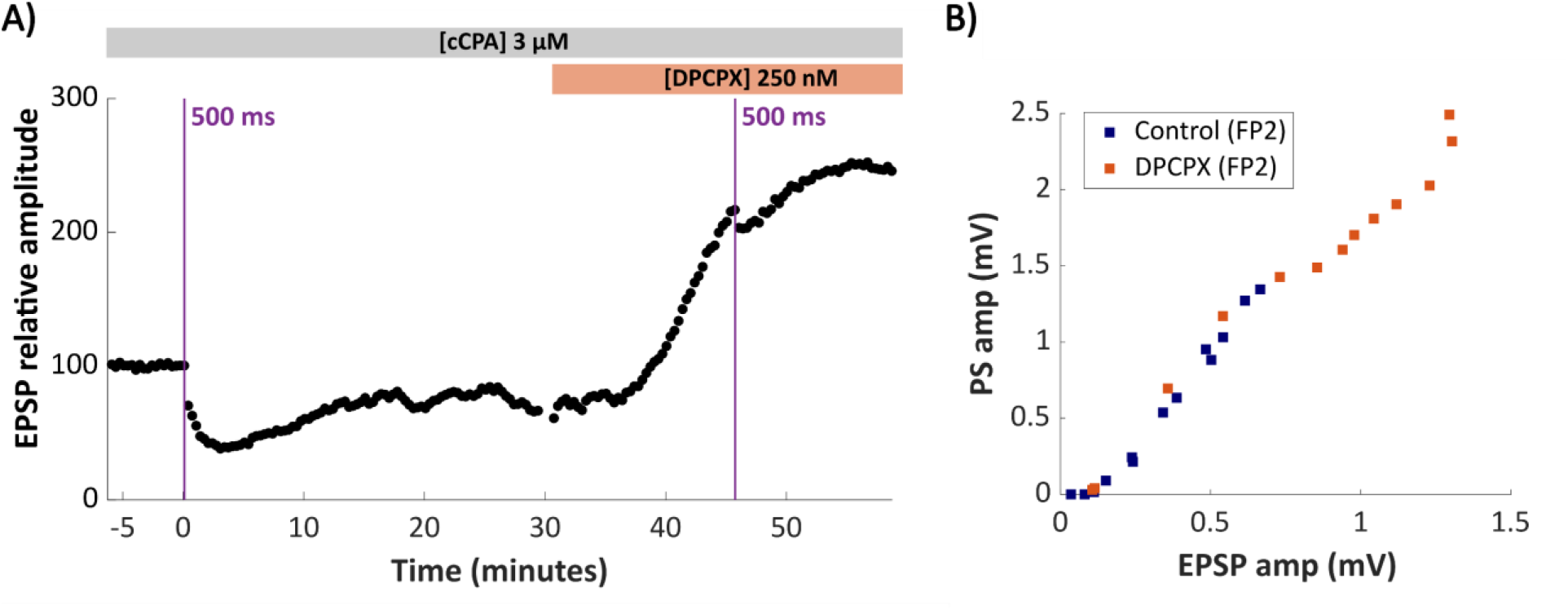
The CPA response is suppressed by the A_1_R antagonist DPCPX. **A)** Time course of the EPSP amplitude from FP1 in the presence of cCPA. After stable baseline recording, a 500 ms flash (t=0) released CPA and transiently reduced the EPSP amplitude. After 30 minutes, DPCPX (250 nM) was added followed by another 500 ms light flash at t=46 mins. **B)** DPCPX increased the EPSP from FP2, reflecting enhanced neurotransmission. Because the relation between PS and EPSP was not changed, it did not change excitability.

These four slices, used for the DPCPX experiments, were part of a larger experiment in 10 hippocampal slices that studied temporal dynamics of excitability changes upon pulsed release of CPA using a single light flash. In slices where at least one channel with a PS in FP2 was observed (9 out of 10), the time course of the EPSP (**Figure 4A**) and of the PS (**Figure 4B**) amplitude were recorded. The CPA pulse reduced the EPSP amplitude in FP2 up to 60.8±4.4% and PS amplitude up to 17.3±8.4% of their baseline values. The reduction was followed by a recovery phase that was considerably slower for the PS than for the EPSP. Superimposing the time course of neuronal excitability (defined as the relation between PS and EPSP) after release of CPA on top of the E-curve constructed from the SR-curves during baseline (as in **Figure 1G**) revealed a clear rightward shift as the system cycles back to its original level (arrows in **Figure 4C**). This shift of the E-curve was variable between experiments, though always present, and implies that a pulse of CPA is followed by a substantial period (minutes) of reduced excitability.

**Figure 4.**
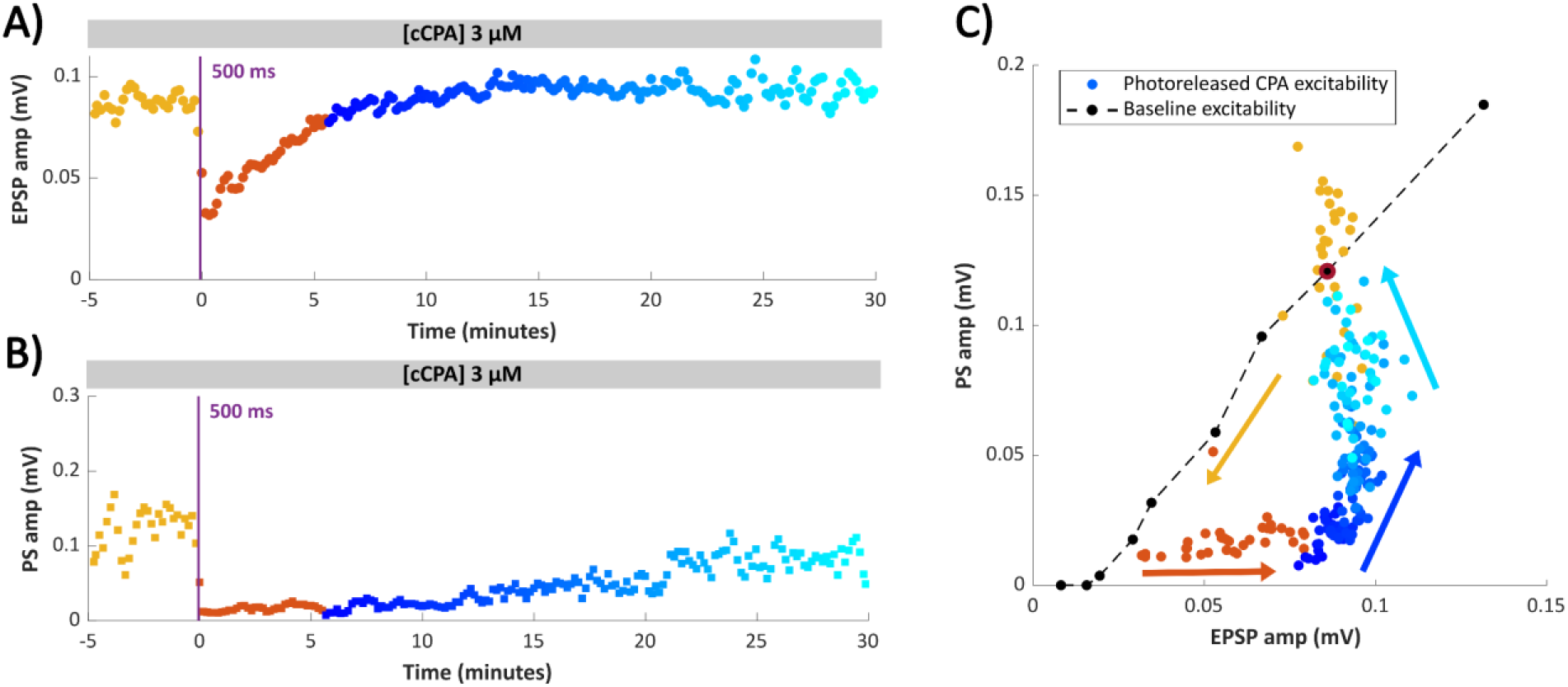
Flash photolysis of CPA with a light flash results in a transient suppression of EPSP and PS. **A)** Time course of EPSP amplitude and **B)** PS amplitude, both from FP2, following a 500 ms light flash that released CPA. Colours indicate the various time periods: baseline (yellow), reduction (orange) and recovery (from dark blue to light blue). **C)** The baseline E-curve, deduced from the SR curves during baseline, for this slice (black dots and dotted line) relates PS amplitude to EPSP amplitude. Superimposed is the excitability time course that the slice goes through after the light flash: identical colour coding for time as used in **(A)** and **(B)**. The arrows in corresponding colours indicate time.

### 3.4 Closed-loop feedback control of neurotransmission with photoreleased CPA

The steep CPA concentration dependence of EPSP suppression (Hill coefficient of 3) implies a narrow therapeutic window, which makes proper dosing for precise changes in neurotransmission and/or excitability challenging. However, its fast activation and slow recovery dynamics make CPA ideally suited for modulation with short pulses such as the ones that result from photolysis of cCPA. To follow up the previous proof-of-principle study [14], the feedback system used to control EPSP amplitude at a user-defined target level was improved and extended. In its simplest implementation the system consisted of an “on-off” regulator where hardware detected level crossing of the FP signal in a user-defined time window and MEA channel. A channel in the Stratum Radiatum that showed a clear dendritic FP with a clear EPSP void of PS, was selected for monitoring. Level crossing triggered a digital pulse of fixed duration (25 ms) that drove the 405 nm light flashes for release of CPA. In **Figure 5** a typical experiment is illustrated where a slice was superfused with 3 *µ*M cCPA and stimuli (intensity I_50%_) were applied to a MEA channel in stratum radiatum every 10 seconds to continuously monitor EPSP amplitude. After ample time for the FPs to stabilize, the reference value (black dotted line) was determined at t=0, the target (orange dotted line) was set to 50% of reference and the loop was closed (indicated as ON). Each time the FP crossed the target level it triggered a light flash with subsequent CPA release (vertical purple lines). In the initiation phase (blue horizontal bar) this happened every stimulus and resulted in a stepwise build-up of CPA that gradually reduced the EPSP amplitude.

**Figure 5.**
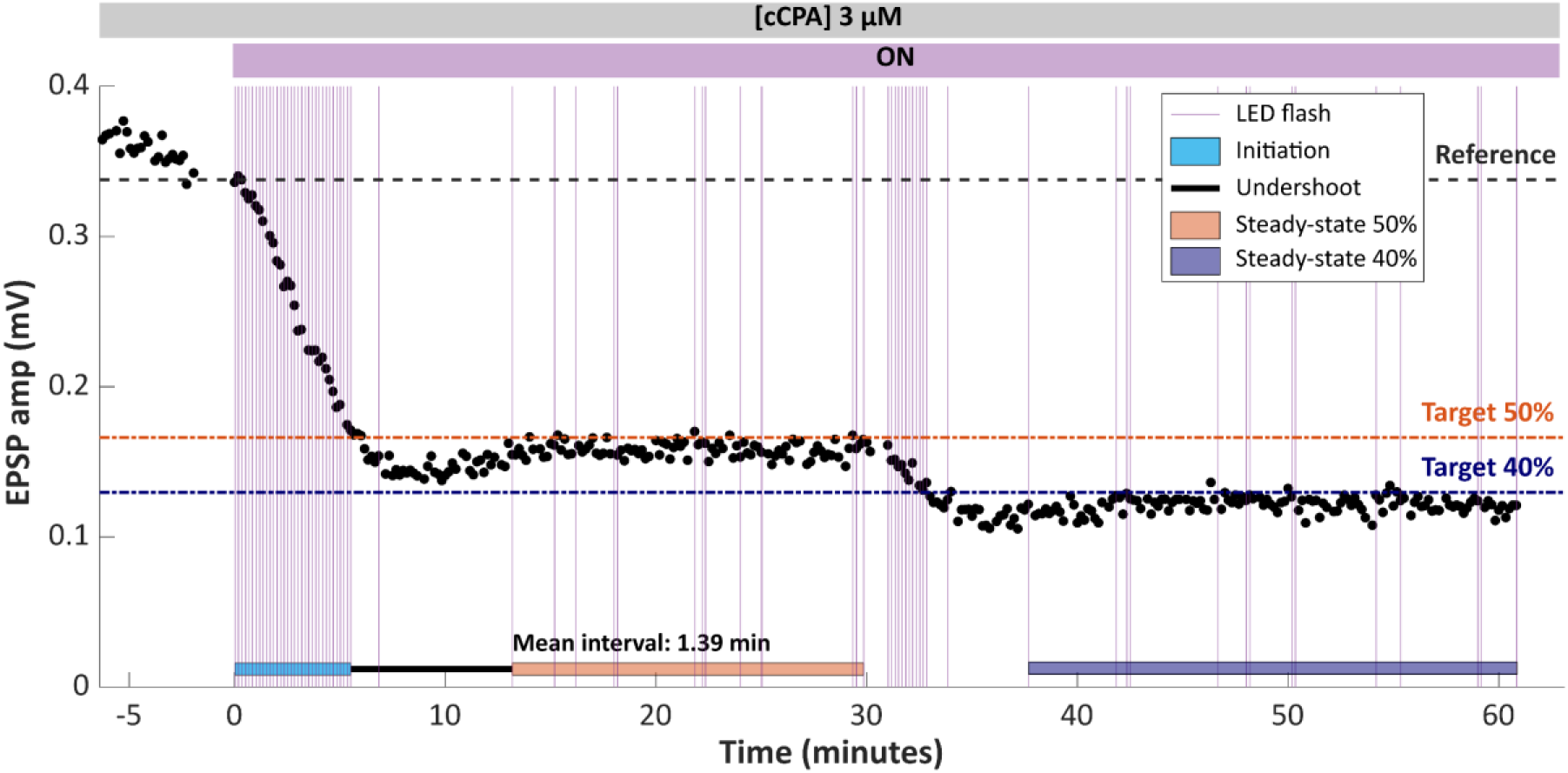
Real-time closed-loop feedback control of EPSP amplitude. FPs were evoked every 10 seconds with I_50%_ stimuli and were allowed to stabilize in the presence of 3 *µ*M cCPA. At timepoint (t=0) the EPSP amplitude was fixed as the reference (black dotted line), the target value was set to 50% of this reference (orange dotted line) and the loop was closed (ON). In this configuration hardware detected level crossing within a pre-set time window after the stimulus and triggered a 25 ms pulse that drove the LED flash. In the initiation phase (light blue bar) each stimulus generated a flash and raised CPA. This gradually brought the amplitude below the target level followed by an undershoot (black bar). Thereafter the system reached steady-state at the target level (orange bar) maintained by flashes with a mean interval of 1.39 mins. At t=30 the target was lowered to 40%, followed by a similar sequence of events : steady-state at the target level (dark blue bar) with a mean light flash interval of 1.58 mins (statistically not different (Mann-Whitney U, p=0.9) from the intervals in the 50% period; the recording period was not long enough to pick-up the expected decrease in light pulse interval. Data points in this figure are offline calculated EPSP amplitudes, while the triggers were hardware generated level-crossings.

This process stopped after reaching the target level and, after several minutes of undershoot (black bar), a stable EPSP amplitude around target was established (steady-state; orange bar). This level was further maintained by light flashes with a mean interval of 1.39 mins. After 30 minutes the target was lowered to 40% and the sequence of events repeated itself: first an initiation phase where every stimulus led to a trigger, followed by an undershoot whereafter again a steady-state situation was reached. The parameters obtained in the [CPA] experiments (**Figure 2**) and (**eq 1.4**), allowed to calculate the CPA concentration needed to keep the 50% target (44.1 nM) while for the 40% target 50.1 nM was needed. Under the simplified assumptions: 1) that the same fixed amount of CPA was released at each light flash and 2) that CPA was removed from the extracellular space by a first-order process, it was also possible to estimate (details in Materials and Methods) the increase in CPA concentration by each light flash (11.9 nM) and the time constant of the washout process (5.2 mins). Depletion of cCPA by a single light flash that generated a 11.9 nM increase in CPA was less than 0.4% and thus negligible. Target settings between 90% and 10% of reference value needed CPA levels between 20 nM and 90 nM (based on **eq 1.4**), which could all be attained with the current settings of the closed-loop.

### 3.5 Elevated potassium increased excitability and induced epileptiform bursting

Functional neurotransmission and stable excitability are crucial for normal brain function and are impaired in epilepsy. The next question was whether the closed-loop system could be used to control, or even prevent, epileptiform bursting while still permitting neurotransmission at an acceptable level? Here, a raise in [K_+_]_o_ was chosen to induce epileptiform bursting. It enhanced excitability and ultimately induced a regular and reproducible form of epileptiform bursting. In a pilot experiment (**Figure 6**) [K^+^]_o_ was raised from a baseline of 3.25mM to 6.5 mM which was calculated to shift the reversal potential for potassium by an estimated 17.5 mV and, based on the Goldman equation [15], to depolarize neurons by 15 mV. FPs were evoked using stimuli of 1250 (I_30%_), 1750 (I_50%_), and 2250 mV (I_70%_), alternating between single and paired pulse stimulation to monitor neurotransmission during the transition over the large excitability range. **Figure 6A** illustrates the time course of the EPSP amplitudes for a dendritic channel. Once the high [K^+^] solution entered the bath, the amplitude of the EPSP gradually increased and reached a higher stable value after 11.4 mins of respectively 257%, 234% and 236% of baseline values (mean EPSP amplitudes measured during the last 15 mins of baseline). After 40 mins a second SR-curve of the EPSP was measured. Using LSSM this SR-curve for FP1 was 2.5±0.1 times larger in elevated potassium than in baseline (weighted mean of the LSSM factors over 21 channels). This factor was even larger than the one obtained for FP2 over FP1 during baseline: 1.8±0.1 (**Figure 6B**, red dotted lines). LSSM was also performed on the time courses of the EPSP amplitudes evoked by the three stimulus intensities during the 6.5mM period. The curves evoked with 1750 and 2250 mV intensity were 1.7 respectively 2.2 times larger than the one evoked with 1250 mV. Remarkably, apart from this scale factor, the shape of the time course was similar (red dotted curves in **Figure 6A** represent the scaled LSSM curves). It is concluded that stimulus intensity is not a critical factor to monitor changes in neurotransmission, all three curves demonstrated similar changes over time.

**Figure 6.**
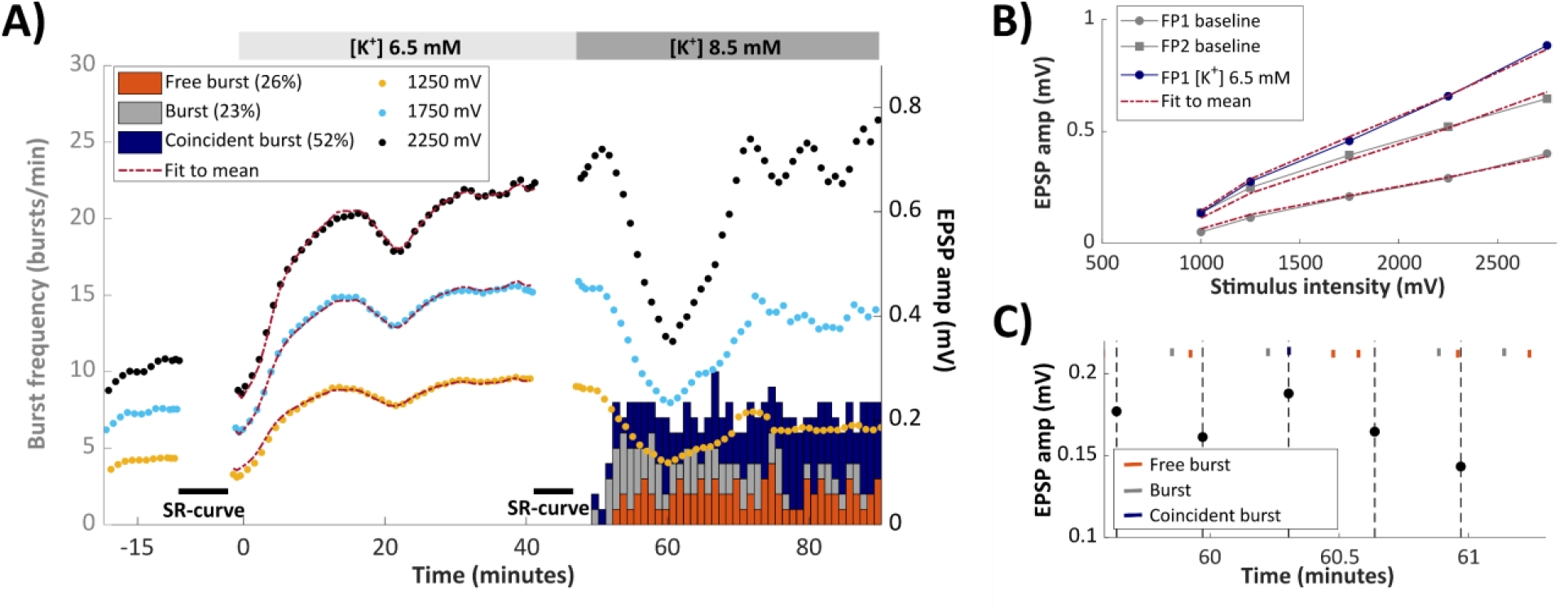
Elevating [K^+^]_o_ increased neurotransmission and induced epileptiform bursting. **A)** Time course of the EPSP amplitude in FP1, evoked with stimulus intensities of 1250, 1750 and 2250 mV, alternating between single and paired pulse stimulation (paired pulse data not shown). At t=0, [K^+^]_o_ was increased from 3.25 to 6.5 mM and at t=45 mins it was further increased to 8.5 mM. Dotted red lines in the period 0-40 mins indicate the LSSM fits of the three traces (factors: 1.0, 1.7 and 2.2). Black bars indicate time periods when SR-curves were recorded (see **(B)**). Histogram bars quantify the burst frequency in bins of 1 minute; colours indicate the three classes (see **(C)**). **B)** SR-curves of the EPSP amplitude of FP1 (grey circles) and FP2 (grey squares) during baseline and of FP1 in 6.5 mM [K^+^]_o_ (blue). The dotted red lines are the LSSM fits to the SR-curves with factors of 1.00, 1.78 and 2.24. **C)** Epileptiform bursts were separated into three classes based on their timing relation with the adjacent stimulation: 1) “free” bursts (orange) were preceded by another burst and thus could not have been directly affected by a previous stimulation 2) “coincident” bursts (blue) occurred within 250 ms after a stimulation and might have been affected by it and 3) all other bursts (grey), which followed a stimulation (0.25–10 sec).

While in the presence of 6.5 mM [K^+^]_o_, none of the MEA channels showed signs of (spontaneous or stimulus-induced) epileptiform bursting, this changed drastically once [K^+^]_o_ was increased to 8.5 mM (at t=45 mins). This second increase in [K^+^]_o_ added an extra (calculated) shift in the K^+^ reversal potential of 6.7 mV, and depolarized the membrane by another estimated 5.7 mV [15]. In this experiment, as well as in all following experiments where [K^+^]_o_ was directly switched from 3.25 to 8.5 mM, the appearance of epileptiform bursts was often accompanied by an unexplained transient dip in EPSP amplitude. Once epileptiform bursting started, it quickly reached a stable frequency of 7.2 bursts/min (**Figure 6A**). FPs and epileptiform bursts and were distinguished by their amplitude and their spatiotemporal voltage profile as illustrated in **Figure 7**. The difference in amplitude and in spatiotemporal morphology allowed to detect spontaneous epileptiform bursts and to easily separate them from the stimulus-evoked FPs. Epileptiform bursts are all-or-none phenomena, once initiated they run their course with almost identical spatiotemporal morphology. This simplified further analysis and allowed to treat them as point processes, characterized by their moment of occurrence.

**Figure 7.**
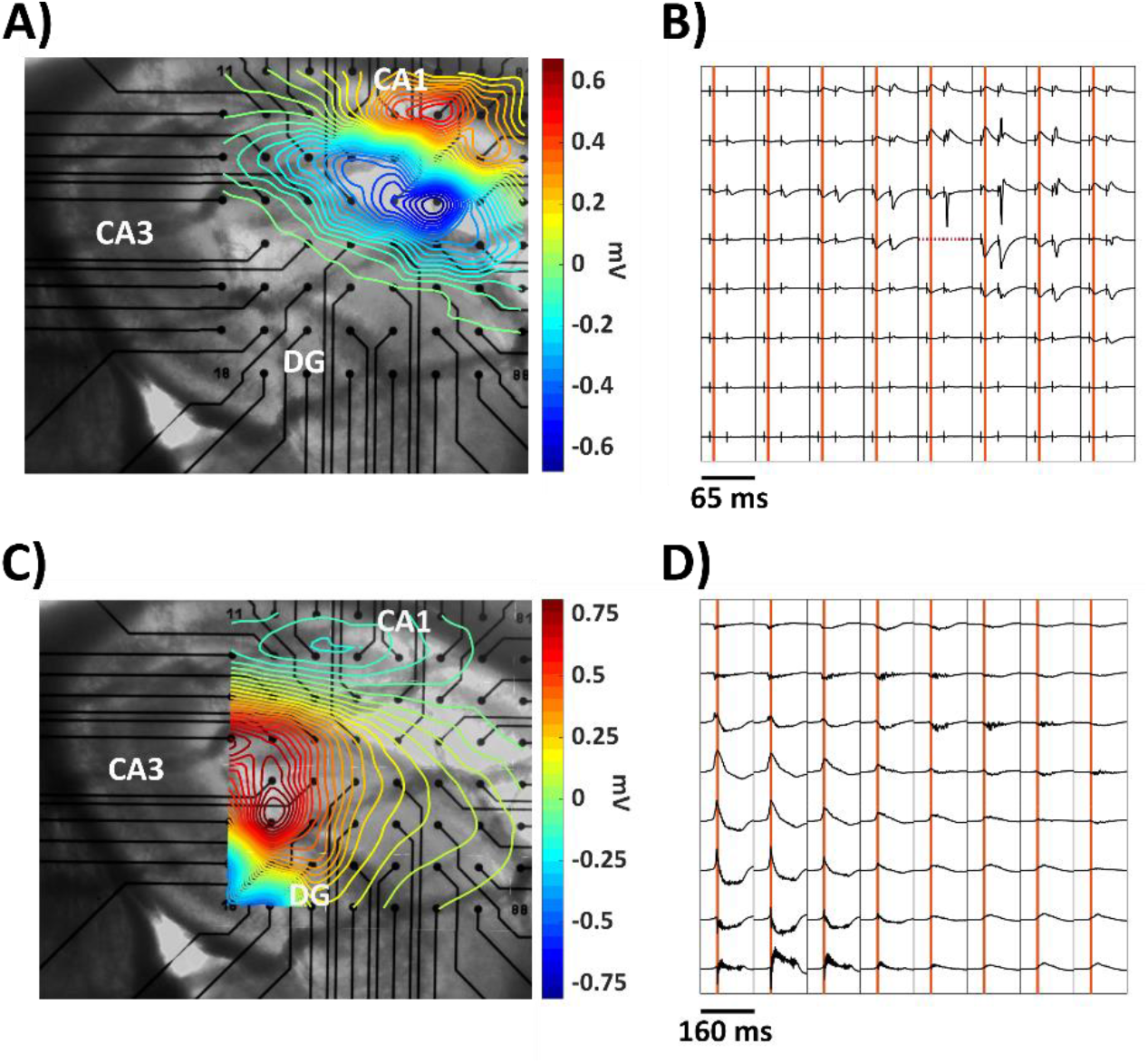
Spatiotemporal voltage profile of a FP and an epileptiform burst in 8.5 mM [K^+^]_o_. **A)** The voltage profile, superimposed on the camera image of the slice, was taken at the time point of maximal EPSP amplitude in the most prominent dendritic channel (vertical orange lines in **B**). **B)** Time course of a typical CA1 paired pulse stimulation at the Schaffer Collaterals. Stimulus channel dotted line. **C)** Similar display as in **A)** for an epileptiform burst in 8.5 mM [K^+^]_o_. The voltage profile was taken at the time point of the earliest maximum (vertical orange lines in **D)**). The burst most likely originated in the CA3 region located outside the MEA grid and propagated towards the other regions. Note the difference in amplitude and in time scale of **B)** and **D)**.

If FPs are to be used for monitoring neurotransmission in a closed-loop system, it is important to know whether there is interference between them and the epileptiform burst, the phenomenon to control. In other words, it had to be checked whether FP stimulation affected timing of epileptiform bursts and whether epileptiform bursts influenced FP amplitude? Based on their moment of occurrence bursts were divided into three classes (**Figure 6C**): 1) bursts that follow another burst were called “free” bursts, as they cannot be directly affected by a stimulation, 2) bursts that closely followed a stimulation (<250 ms) but had not started before it, could have been affected by it and were called “coincident” bursts and 3) all other bursts. The three classes did not differ in spatiotemporal voltage profile: they only differed in their time relation to the stimulus. The relation between FP stimulation and the timing of bursts in the experiment illustrated in **Figure 6A** is given for the three stimulus intensities for single (**Supplementary Figure S1A**) as well as paired pulse stimulation (**Supplementary Figure S1B**). For regular bursting, chance predicts that about 2.5% of the bursts fall in the first 250 ms of the 10 s interval that separated two stimuli. In this initial experiment the fraction of those coincident bursts was considerably higher. It increased with stimulus intensity (40.8%, 41.7% and 50.9%), and with the same intensities paired pulse stimulation gave more coincident bursts (47.5%, 61.4% and 63.8%). It was not clear whether the stimulation only changed threshold or directly triggered the burst, but the results indicated that to minimize the chance of interaction between FP and bursts it was necessary to stimulate with intensities considerably lower than 1250 mV (I_30%_).

Therefore, in all following experiments, monitoring stimulus intensity was set to I_10%_ while continuously recording for the presence of epileptiform bursts upon switching [K^+^]_o_ from 3.25 to 8.5 mM. An example experiment, displayed in **Figure 8**, shows that EPSP started to gradually increase almost immediately after the concentration switch and reached a peak value of 272% of baseline after 22.7mins (**Figure 8A**). The first epileptiform burst was detected 11.6 mins after the switch when the EPSP amplitude had risen to 160%. The burst frequency quickly stabilized and reached after 45mins a mean value of 4.6 bursts/min. In **Figure 8B** a somatic and a dendritic channel were chosen to illustrate the development of the FPs at three distinct timepoints during the transition. The EPSP of FP1 increased in amplitude with increase in [K^+^]_o_, but the I_10%_ stimuli did not induce multiple PSs. FP2 was larger and developed a few PSs after 5 mins (yellow trace) and multiple PSs after 15 mins when the epileptic state was reached.

**Figure 8.**
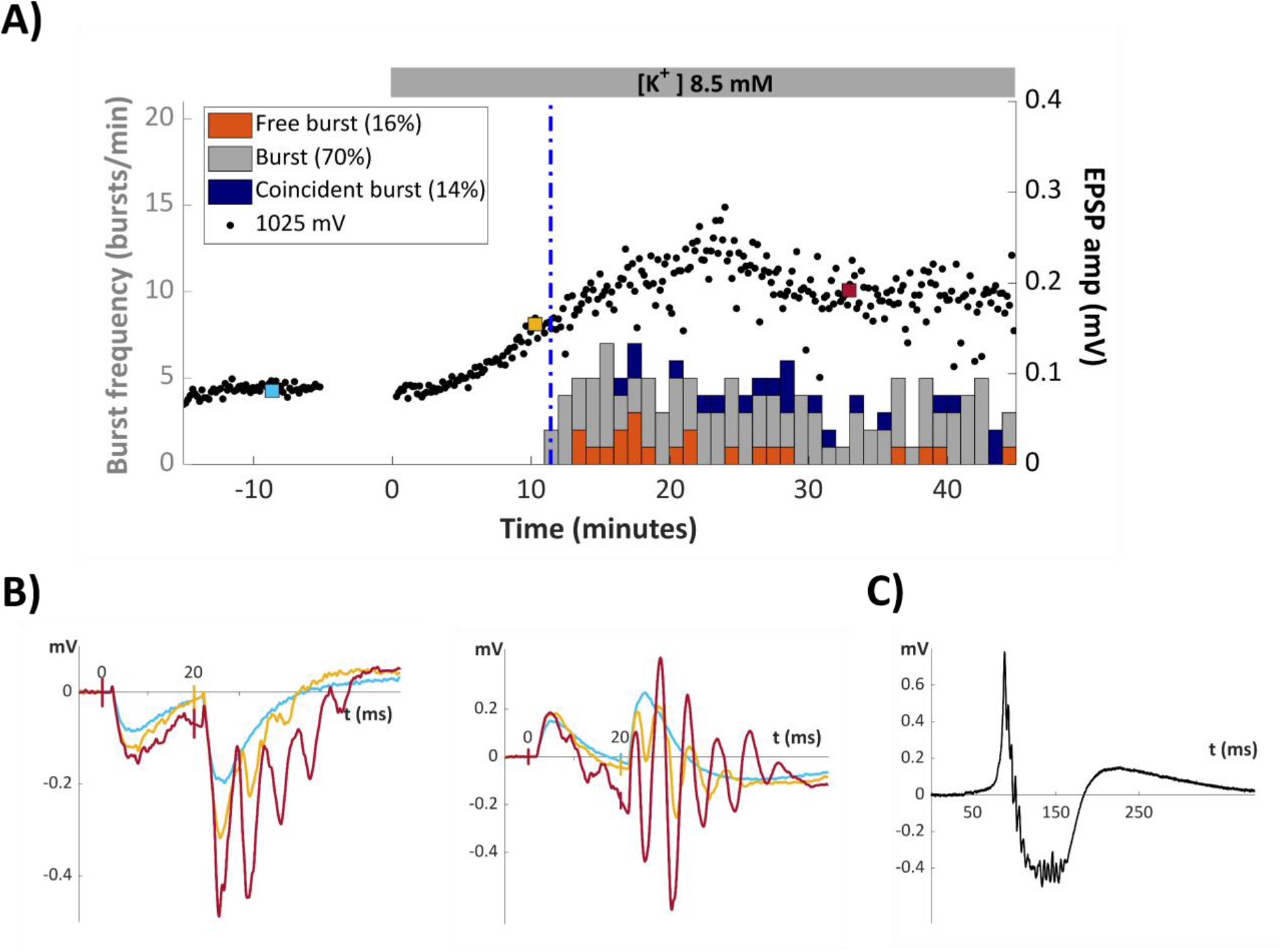
FP evolution induced by [K^+^]_o_ transition from 3.25 to 8.5 mM. **A)** Time course of the EPSP amplitude evoked with a stimulus intensity of 1025 mV (I_10%_), alternating between single and paired pulse stimulation (only FP1 data shown). After 15 mins of baseline recording (at t=0) [K^+^]_o_was increased from 3.25 to 8.5 mM. The EPSP started to increase in amplitude and reached a peak value of 272% after 22.7 mins. The first burst was detected 11.6 mins after switching to 8.5 mM (vertical blue line). At that timepoint EPSP amplitude had risen to 160% of its baseline value. Burst frequency stabilised and reached a stable frequency of 4.6 bursts/min after 45 mins. **B)** dendritic negative EPSP (left panel) and somatic positive EPSP (right panel). They are presented at three different timepoints: square markers in **A**: baseline (blue), initial rise (yellow) and near the end when EPSPs were stabilized (red). **C)** Typical epileptiform burst from an electrode near the CA3 region.

**Figure 8C** illustrates a burst recorded from an electrode located near the CA3 region; this signal is twice as large and lasts approximately 10 times longer than the EPSP. **Figure 9A illustrates** the effect of FP stimulation for single (top panel) and paired (bottom panel) pulse stimulation with I_10%_ intensity. For single pulses, the distribution of the bursts over the 10 seconds interval between the stimuli was uniform and the fraction of coincident bursts (2.9%) not different from what was expected from chance. For paired pulses, which in elevated potassium produced multiple PSs, a larger fraction of coincident bursts (13.5%) was detected, suggesting mild interaction. This did not seem sufficient to drive/synchronize the bursting pattern, because the histogram was uniform over the 0.25-10 seconds time interval. In **Figure 9B** the inverse relationship was investigated showing that a preceding burst did not affect the EPSP amplitude in a time dependent manner. The aggregated result for this series of experiments (*n=3*) was that in all cases elevating [K^+^]_o_ to 8.5 mM induced bursting which reached a stable mean frequency of 7±1.7 bursts/min, the first burst appeared 8.4±1.7 mins after the switch at a time point where the EPSP had increased to 166±23% of baseline.

**Figure 9.**
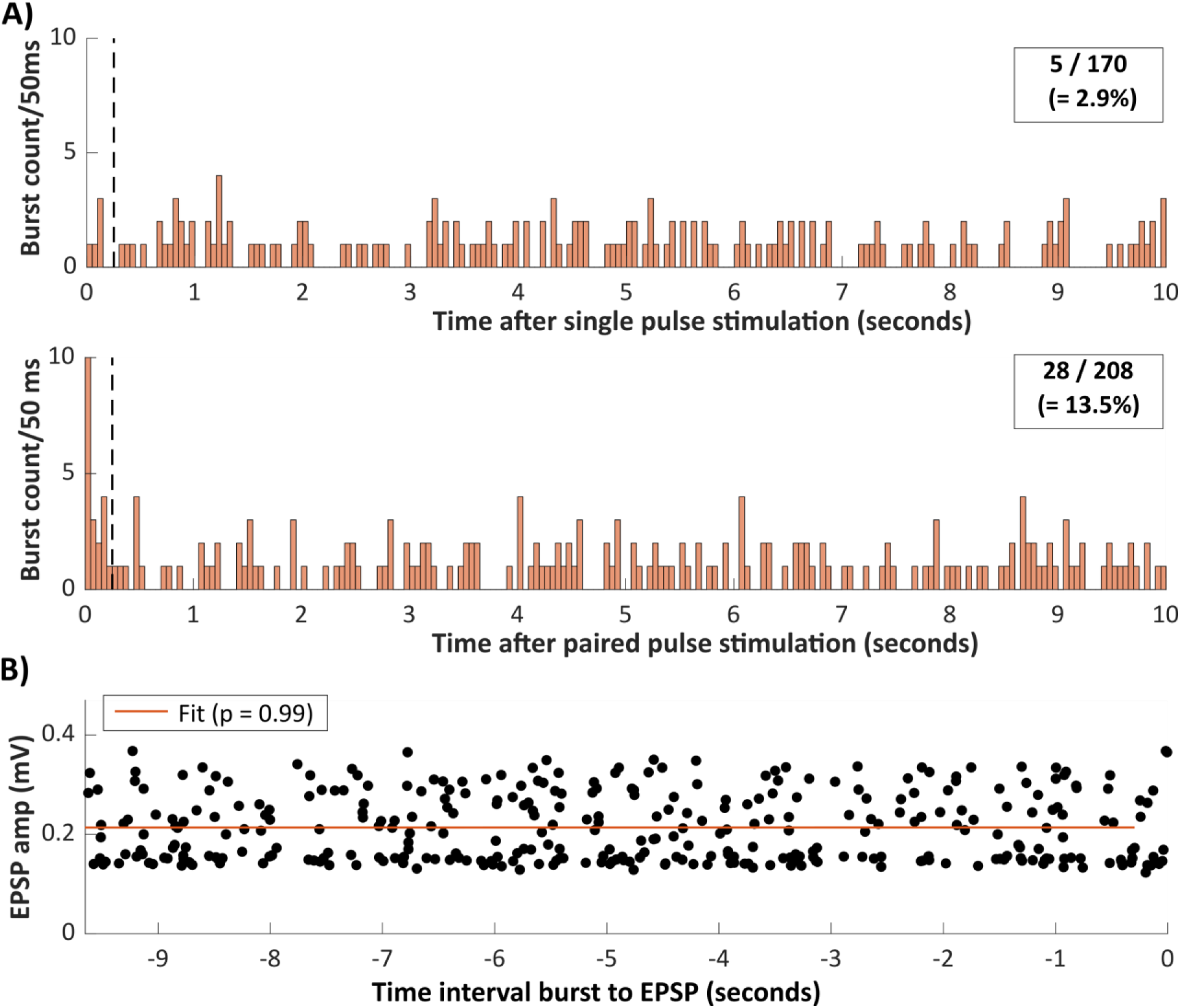
Interaction between FPs and epileptiform bursts. **A)** Likelihood of a burst to occur at a certain moment after a single (top panel) or a paired pulse stimulus (bottom panel) expressed in counts/50 ms. Coincident bursts fall within the first 250 ms of the histogram and make up 2.9% of the bursts observed after single pulse stimulation; a fraction that is not different from what was expected based on chance. After paired pulse stimulation this fraction was larger (13.5%), suggesting that at least some of the bursts were facilitated by the paired pulse. This did not seem sufficient to drive/synchronize the bursting pattern. **B)** Every datapoint at time t in this graph represents the EPSP amplitude of a FP at time 0 that is preceded by a burst at time t. There is no relation between the amplitude of the EPSP and time-interval between burst and EPSP.

### 3.6 Controlling neurotransmission and epileptiform bursting with feedback photorelease of CPA in elevated potassium conditions

In the following experiments the closed-loop control system was modified. Instead of using hardware level crossing to trigger a LED pulse of fixed duration, two (selectable) MEA electrodes were continuously monitored with an external ADC. One recorded an optimal dendritic FP response, the second one a somatic FP response that at higher intensities also displayed a PS. Real-time analysis of these two FPs continuously provided all parameters (EPSP amplitude, PS amplitude and CLI) that were put in an algorithm that generated the light flashes and thus controlled CPA release.

The switch from 3.25 to 8.5 mM [K^+^]_o_ resulted in a first epileptiform burst after 8.4±1.7 mins when the EPSP amplitude was 166±23% of its original value. The next trials investigated whether preventing the increase in EPSP (partially or completely) with feedback-controlled photorelease of CPA was sufficient to prevent the appearance of epileptiform bursts. A similar strategy as in the previously described closed-loop experiments was used. Slices were superfused with 3 *µ*M cCPA while alternating single and paired pulse stimuli (I_10%_) were applied to continuously monitor EPSP amplitude. At t=0, the EPSP reference amplitude was determined, and the target level was set to 140% (*n=1*), 120% (*n=5*) or 110% (*n=2*) of this value. Then [K^+^]_o_ was switched to 8.5mM and the loop was closed (indicated by ON1). During ON1 the FPs were real-time analysed and their EPSP amplitudes were compared with the target level; if the EPSP amplitude was larger than target, the algorithm generated a light flash of 50 ms that released CPA.

**Figure 10A** gives an illustrative example for a target of 110%. After the switch to 8.5 mM [K^+^]_o_ the EPSPs gradually increased and as soon as their amplitude reached 110% the loop became active (light flashes are indicated by vertical purple lines). Sufficient CPA was released to prevent further increase of the EPSP amplitude, and in this experiment, epileptiform bursts were prevented for at least 30 mins.

**Figure 10.**
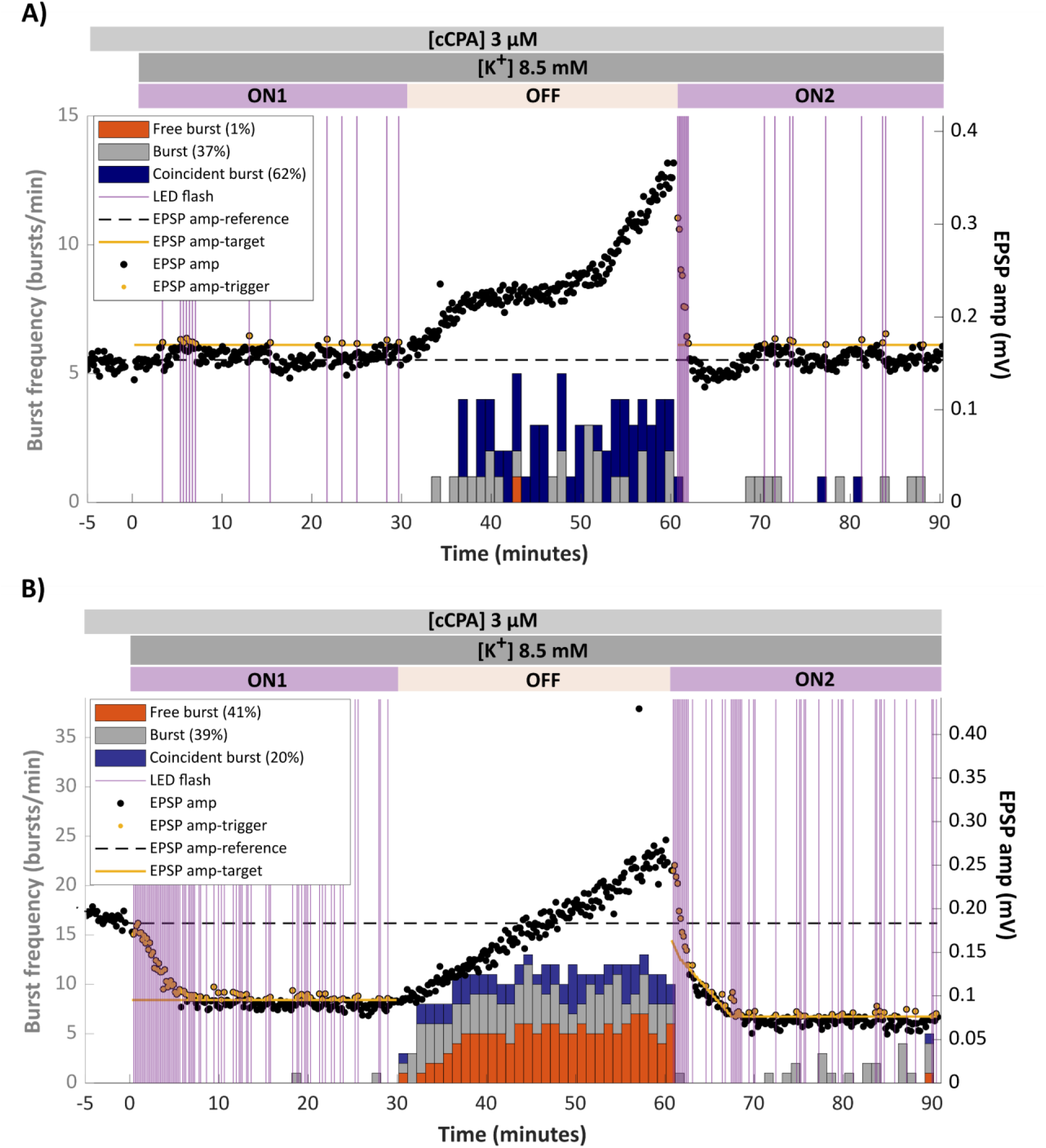
Closed-loop control of EPSP amplitude in elevated [K^+^]_o_ suppresses epileptiform bursting. Time course of the EPSP amplitude (FP1 response) in the presence of cCPA. At time t=0 reference amplitude was determined, target amplitude was set to either 110% in (**A**) or 50% in (**B**), [K^+^]_o_ was raised to 8.5 mM and the feedback loop was closed (ON1). During this period EPSP amplitudes that surpassed target level (yellow line) generated a 50 ms light flash (vertical purple lines) that released CPA and stabilized EPSP amplitude around target level. After 30 mins, the loop was opened for 30 mins (OFF) which resulted in a gradual increase of EPSP amplitude and the appearance of bursts. At 60 mins, the loop was closed again (ON2), which quickly brought back EPSP amplitude to target level and strongly suppressed bursting.

In half of the experiments, a temporal dip in EPSP amplitude was observed about 10 minutes after the switch to 8.5 mM [K^+^]_o_ (see also **Figure 6A**). This dip was not accompanied by a (transient) change in the bursting rate. For a closed-loop system the relation between the control variable (EPSP amplitude) and the controlled phenomenon (epileptiform burst) need not to be known and may be (partially) nonlinear, but it must be monotonous for effective control. Although it was never observed, such a dip could lead to (temporally) loss of control by the feedback loop.

Although sometimes very effective, bursts were not always suppressed at target levels above 100% and in an additional set of slice experiments, target level was lowered to 50%. Such an experiment is illustrated in **Figure 10B** and despite the fast increase in CPA and associated decrease in EPSP in the first 5 minutes the time course in ON1 is very similar to the one in **Figure 10A**. As was expected the relative burst suppression with target at 50% (89±6%, *n=5*) was slightly higher than that for the three previous target settings (80±8%, *n=8*), but the difference did not reach significance and might well be irrelevant.

As a positive control the loop was opened after 30 minutes (indicated by OFF, **Figure 10A**) whereafter the EPSP amplitude increased and after 5-10mins epileptiform bursts appeared in all experiments. Burst frequency stabilized after 10-15 mins in the OFF period to a value (8.2±1.3 bursts/min, *n=13*) that was not different from the frequency measured in the elevated [K^+^]_o_ slices not exposed to CPA (7±1.7 bursts/min). Sixty minutes after the switch to 8.5 mM, the loop was closed again (period ON2) to validate that it was still able to reduce the EPSP amplitude to target level. This substantially, but not completely, suppressed the epileptiform bursts (**Figure 10A**).

Averaged over all experiments burst suppression in the ON1 phase (83±5%, p<0.001), as well as in the ON2 phase (79±5%, p<0.001) was very effective and the two phases were not statistically different.

In all experiments so far, the closed-loop monitored and controlled the EPSP amplitude and thus specifically neurotransmission and not excitability. But epilepsy and epileptiform bursting are more a problem of excitability than of neurotransmission. The EPSP amplitude varied over a wide range, and the experiments showed that the feedback loop can modulate it over almost that full range. In contrast the transition from stable baseline to bursting seemed to be more of an abrupt transition and burst frequency did not gradually follow EPSP amplitude over its full range. The optimal control solution needs to find a target level that maximizes the range of neurotransmission, while excitability is just below the threshold for bursting. Preferentially, the closed-loop system should be able to find that level in an unsupervised way.

The experiments illustrated in **Figures 5, 7 and** **9** demonstrated that the increase in PS amplitude as a function of time was always larger than that of the EPSP, implying that excitability substantially increased in elevated potassium conditions. **Figure 8** shows that ultimately stimulation led to multiple PSs. The CLI is a measure that elegantly combines PS amplitude and the presence of multiple PSs into a single number and is therefore a strong measure for epileptogenicity. The paired pulse stimulus often used in this study also came in extremely useful, because, for the right stimulus intensity, FP1 offered an optimal EPSP while FP2, where excitability was substantially higher, presented multiple PSs once excitability increased (**Figure 8B and Figure 11**). In the last series of experiments, the CLI was added to the decision algorithm that generated the light flashes. It was calculated from the second external MEA signal that provided the somatic FP. As before the light flash was generated when the EPSP amplitude was larger than the target level, but now the target level was put under control of the CLI. The CLI reference value was set at the same moment (t=0) as the target reference and a maximum value for the CLI was arbitrarily defined (120%); the control algorithm compared the CLI with the chosen maximum value and when it was larger the algorithm reduced the EPSP target level by a user defined step (2%).

**Figure 11.**
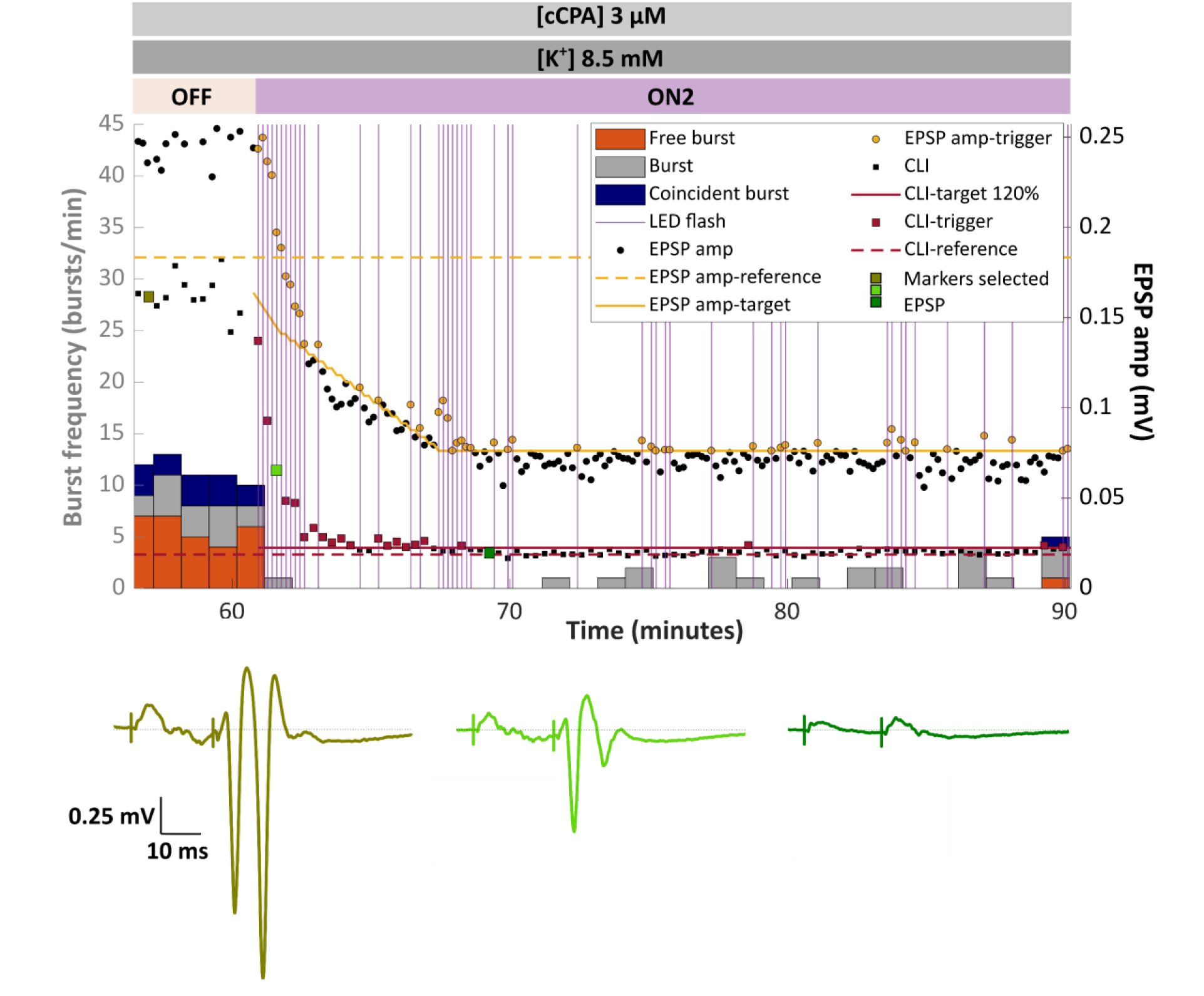
Closed-loop control with a dynamic target linked to excitability (CLI). Control of dendritic EPSP amplitude and bursting in the ON2 phase in the system that was extended with a dynamically adjusted target level (yellow solid line). Light flashes and CPA release were generated, as before, only when EPSP amplitude in FP1 was larger than target level. Target level initially started at 90% of its reference value (yellow dotted line) and was downward adjusted by 2% at every somatic FP where CLI (in arbitrary units, dominant in FP2) was larger than a user-defined maximum (120%, red solid line). Around 68 minutes the target level hit its pre-set minimum of 40%. Lower panels show three paired pulse stimuli evoked at moments in time indicated by green square markers in the CLI trace; EPSP was initially much larger than 100% to be finally reduced to 40% of the reference value. PS and CLI (best observed in FP2) were reduced even more strongly.

This extended mode of operation was implemented, and its behaviour was investigated in the ON2 period, the phase of the experiment that started from a condition where stimuli generated multiple PSs, and bursts were present (**Figure 11**). By the end of the OFF phase the CLI, expressed in arbitrary units, had reached about 500% of its baseline value and thus way above the chosen maximum. The initial EPSP amplitude target level was set at a relative high value (90%), and the decremental step-size for the target level was fixed at 2%. Also, a (selectable) lower limit of 40% for the EPSP target was introduced to prevent complete shutdown of neurotransmission. **Figure 11** presents all relevant signals for the system in the ON2 phase of the experiment depicted in **Figure 10B**. By the end of the OFF period the EPSP amplitude as well as CLI were both much larger than their respective targets (yellow solid line and red solid line). This implied that when the loop was closed again (ON2), the algorithm generated a flash for each FP, and it also reduced the target level by the defined step. The photoreleased CPA started to decrease EPSP and CLI in a way that it reduced excitability. Once the CLI dropped below the chosen maximum level (120%), the EPSP target level was kept stable, but light flashes to maintain the EPSP amplitude below that target were still needed. In this specific example target just hit its lower limit (40%), and not all bursts were suppressed; a slightly lower limit would have been preferred. In the bottom panel (**Figure 11**) three samples of the paired pulse somatic FPs, taken at relevant moments in time (green markers) illustrate the effectiveness of the new configuration. Note that FP1 provided a clear EPSP signal while FP2 provided a signal with multiple PSs.

The expanded system has three additional parameters (CLI maximum, EPSP target step size, EPSP target lower limit), that allowed fine-tuning, which was not performed in any detail at this stage of the experiments. As in all closed-loop feedback systems, these parameters in addition to flash pulse duration and more advanced control loop strategies (e.g., PID control) could be used to optimize speed, improve accuracy and guarantee stability, strongly dependent on the properties of the system to be controlled. These experiments did, however, prove the validity of the closed-loop photocontrolled strategy and are open for further improvement.

## 4. Discussion

Functional neurotransmission and stable excitability are crucial for brain function. They are highly relevant for many brain diseases including epilepsy. In this study a clear distinction is made between 1) neurotransmission: how much synaptic signal (EPSP) is obtained on activation of a certain fibre tract and 2) excitability: how much PS is obtained by a certain level of synaptic activity. Despite its complexity, this is a gross simplification of any neuronal region that mostly involves feedforward and recurrent networks of excitatory and inhibitory neurons and many forms of modulation. The goal in epilepsy is to have a therapy that controls excitability below the threshold for seizures while still maintaining a defined level of neurotransmission, necessary for brain function and to avoid adverse side effects. The A_1_R is an ubiquitously present G protein-coupled receptor which, once activated, reduces synaptic transmission and increases firing threshold via inhibition of VGCC and activation of GIRK channels [2]– [4]. Together these two mechanisms create an ideal mix to reduce excitability.

In this study, the non-metabolized A_1_R agonist CPA was used to activate A_1_R signalling. First, the CPA sensitivity of the EPSP was determined in the rat hippocampal CA1 region at the population level under the assumption that the generation rate of active G protein subunits is determined by the occupancy of the A_1_R, while their dissociation to inactive forms occurs with a fixed rate. The observed third order binding properties (Hill coefficient 3) imply that there is only a narrow therapeutic window for A_1_R medication: as an example, the difference between 20% and 80% effect is from 28 to 70 nM, while for a first order kinetics that range would be 10 to 176 nM. However, the fast activation (below the time resolution of our measurements, so at least in the range of seconds) and the slow recovery (time constant around 22 mins, determined in Craey et al. [14]), make the A_1_Rs ideally suited for pulsed application and modulation of the CPA concentration.

Recently, a coumarin-caged form of CPA (cCPA) was developed, which can be uncaged with LED light flashes of 405 nm at millisecond timescale [14]. An additional advantage of the photopharmacology approach is that it gives precise control over where CPA is released, which, given the ubiquitous presence of A_1_Rs in the body, will avoid many adverse side effects. In the current slice experiments uniform illumination was applied and it is assumed this results in a uniform CPA concentration increase in the slice. CPA is a small molecule and we assume its concentration will equilibrate within minutes, a time scale relevant in these experiments. No effects attributable to unequal CPA distribution were ever observed in the slice experiments. This will be much more complicated in a three-dimensional *in vivo* situation.

The time course of EPSP and of PS amplitude after a short pulse of CPA demonstrated that the neuron cycled through a phase of reduced excitability before returning to its stable state. Altogether, these properties led to the design of a simple on-off pulsed feedback system that used information from regularly sampled FPs (EPSP and CLI) to control the level of neurotransmission and excitability. Using fixed pulses that raised CPA levels by about 10 nM, applied by the closed-loop system when the EPSP amplitude was larger than a user-defined target, allowed to control EPSP/neurotransmission over the full 10-90% range of the baseline situation. There are many ways to further optimize this control system. Adaptive light flash duration could optimize the duty cycle of CPA application to bring variation close to the inherent variation in the EPSPs. Tuning flash size proportional to the deviation in EPSP will speed up the transitions. Alsomore optimizations in the direction of PID control are possible, certainly in the extended software implementation of the system, where a user-definable algorithm runs the control loop.

In this study the priority was to investigate in an experimental model of epilepsy whether closed-loop control using cCPA/CPA was able to suppress epileptiform bursting while maintaining neurotransmission. Enhancing [K^+^]_o_ to 8.5 mM was selected as the model that reliably generates epileptiform bursting, not via pharmacological blocking of a specific receptor, but via a mechanism that is involved in many forms of seizure generation [16]–[18]. However, suppressing epileptiform bursts is only a first step towards control of clinical epileptic seizures and one may wonder whether a model based on the reduction in driving force for potassium can be effectively counteracted by a mechanism that relies on enhanced potassium conductance. Therefore, instead of fine-tuning the current system, the next important and relevant step will be to test whether this approach can be generalized to other epilepsy models, including chronic and *in vivo* models and certainly to pharmaco-resistant models. Monitoring the dendritic EPSP gave almost complete control over neurotransmission, but it was clear that to control epileptic activity, excitability is essential and somatic information about bursting activity (multiple PSs) would be very useful. Multielectrode recording with the MEA device permitted to select dendritic channels that provided a clean EPSP. In addition somatic channels allowed quantification of (multiple) PSs with the CLI (=total line length) that included PS amplitude and the count of PSs in a single number. The paired pulse stimulation had the advantage, that in FP2 excitability is considerably higher than in FP1, which makes FP1 optimal for EPSP amplitude and FP2 optimal for CLI determination if the correct (low!) stimulus intensity is used.

Closed-loop systems can run into serious problems if the control variable and the controlled phenomenon interfere. In the current system that could happen if electrical stimuli affect (timing of) bursts, or if bursts strongly influence EPSP amplitude. As long as stimulus intensity is I10% or lower the fraction of coincident bursts with single pulse stimulation was as expected by chance, but for paired pulse it was higher. Indications that stimulation drove, or synchronized bursting were never observed. Also, coincident bursts were not the first ones to appear or the last ones to disappear. Finally, in the stable situation bursts were almost completely suppressed, so that interference automatically disappeared. For the reciprocal question, do bursts preceding a FP affect its amplitude, the situation is even simpler: only bursts that really overlap with a FP corrupt its analysis. Bursts start so slow that for most coincident bursts the FP can still be analysed; less than 2% of the FP are too corrupted to analyse and could be easily skipped by the software of the feedback algorithm. Time-dependent effects of bursts on EPSP amplitude were not observed. Finally bursts and FPs originated in distinct regions of the slice which might limit their interaction to superposition of the electrical fields with relatively little functional interaction.

In recent years, the field of neuromodulation for epilepsy has experienced a significant shift towards the development and application of closed-loop strategies. Until now most consisted of responsive optogenetics and responsive neurostimulation (NeuroPace). Those technologies all use a form of early seizure detection and then try to interfere with the seizure by either optical activation of genetically inserted channels or by neurostimulation [19]–[23]. Several problems are encountered: these methods need a reliable seizure prediction method whereafter the seizure can be prevented instead of aborted. Until now the prediction techniques are rare, and one might wonder whether seizures can be predicted at all. For the methods that need detection there is also a fundamental inconsistency in requiring the presence of seizures in order to prevent them. And finally it seems much more difficult to abort a seizure than to prevent one [24]–[26]. Preclinical approaches that use FP related information about excitability not predicting seizures but estimating their likelihood have been proposed [27] but never implemented.

Our earlier chemogenetic studies [28], [29] demonstrated the powerful seizure-suppressive effects of inducing neuronal Gi signalling specifically in the seizure focus. The current pioneering closed-loop photopharmacological intervention is unique in that it uses the highly specific molecular properties of the adenosine signalling pathway, which directly regulates neurotransmission and excitability. This is a new approach to suppress seizures and in line with the fact that adenosine acts as an endogenous anticonvulsant because the increase in concentration observed during seizures counteracts ongoing seizure activity [1]. Most anti-seizure medication consists of therapeutics that restrict high frequency firing by slowing down removal of sodium channel inactivation. These therapeutics are, however, not successful in around 30% of the patients, so there is an urgent need for other approaches that take fundamentally different routes. Whether treatment with photoreleased CPA could partially fill the gap, needs to be investigated and certainly a lot of practical problems must be resolved before light uncaging can be routinely applied in patients.

The experiments presented in this study, demonstrate that it is possible to design and implement a photopharmacological closed-loop system that uses the A_1_Rs to suppress epileptiform bursts, while preserving neurotransmission at a controllable and reasonable functional level. The control algorithm uses information extracted from dendritic and somatic FPs that monitor the excitable state of the slice preparation but can be expanded to any desired level, which could make the system applicable in other brain diseases that deal with compromised excitability.

## Funding

This work was supported by the Ghent University Special Research Fund, the Queen Elisabeth Medical Foundation, the Margaret Olivia Knip Foundation and Research Foundation Flanders-FWO (grant numbers 1S65521N, G042219N, G075224N).

---

**Figure S1.**
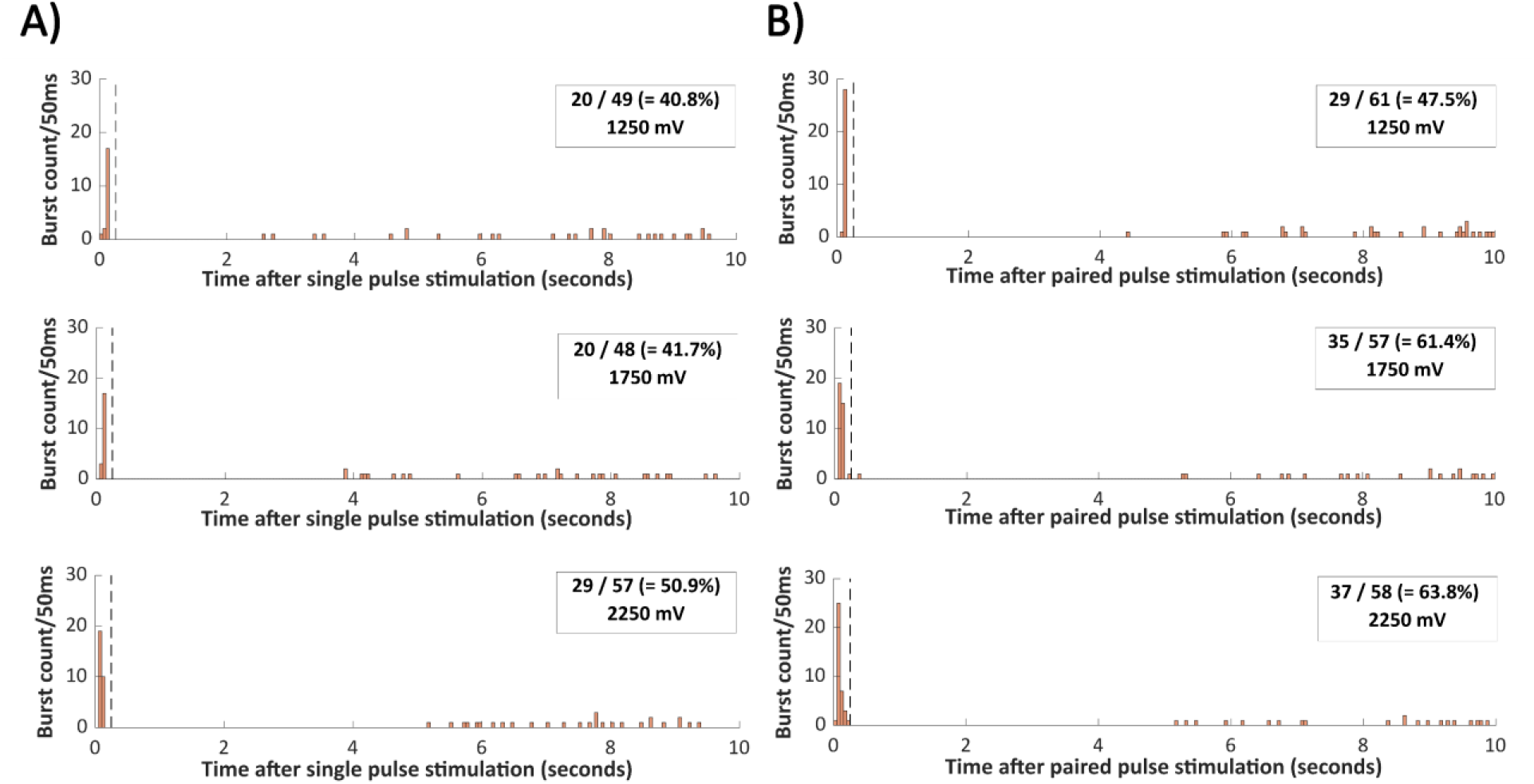
Strong stimulations trigger epileptiform bursts in elevated K^+^ medium. Burst occurrence in the interval between two stimuli, for single **(A)** and paired pulse **(B)** stimulation with three intensities (top to bottom: 1250, 1750 and 2250 mV). Bars represent burst counts in 50 ms bins. The likelihood of finding a burst within the first 250 ms of the histogram (indicated with the dotted vertical line) is much larger than expected by chance. It increases with stimulus intensity and is higher for paired pulse than for single pulse stimulation.

## Notes

### Competing Interest Statement

The authors have declared no competing interest.

